# Individual differences in punished alcohol self-administration are unaltered by alcohol vapor exposure

**DOI:** 10.1101/2025.01.22.634361

**Authors:** Maya N Bluitt, Ana C Munoz, Joyce Besheer

## Abstract

Continued alcohol use despite negative consequences is a defining feature of alcohol use disorder (AUD). It remains poorly understood whether individual variability in drinking despite negative consequences is due to inherent differences or emerges after prolonged alcohol use. The goal of the present study was to use a rat model of drinking despite negative consequences to assess individual differences in foot shock-punished alcohol self-administration prior to and following alcohol vapor exposure in male Wistar rats. After baseline operant self-administration was established, rats underwent additional self-administration sessions in which random, response-contingent foot shock punishment was introduced. Average percent change from baseline was calculated for each rat during punished sessions and rats were classified into shock-sensitive (SS) and shock-resistant (SR) subgroups using the top and bottom thirds. Rats then underwent 3 cycles of air or alcohol vapor exposure every other week, with unpunished self-administration sessions occurring during the intervening weeks. Following the last vapor cycle, rats were re-assessed for resistance to foot shock during punished self-administration sessions. Alcohol vapor exposure had no effect on punished self-administration overall, nor by subgroup. Examination of individual differences showed that rats classified as SR showed increased unpunished self-administration relative to baseline regardless of air vs. vapor condition. These data suggest that alcohol history has a minimal effect on individual differences in foot shock-punished self-administration.

## Introduction

Alcohol use that persists despite negative consequences is a cardinal feature of alcohol use disorder (AUD). Understanding the development and maintenance of this behavior has become a major focus of preclinical research [1, 2, 3]. In rodents, alcohol drinking despite adverse outcomes is often modeled by adulteration of alcohol with a bitter tastant (primarily quinine) or pairing alcohol seeking or consumption with foot shock punishment [2]. While such adverse consequences typically result in attenuation of alcohol drinking, factors such as motivation for alcohol [4], stress [5], sensitivity to negative feedback [6], and access schedules (i.e., continuous vs. intermittent; 7, 8, 9) influence the likelihood of an animal to demonstrate drinking despite negative consequences. However, the extent to which alcohol history influences punished drinking remains unclear.

It is generally accepted that some alcohol history is required to induce alcohol drinking despite negative consequences [1, 2, 10]. For example, rodents exhibit resistance to quinine adulteration after extended access to alcohol (i.e., ≥ 3 months; 7, 11, 12) or chronic intermittent alcohol exposure (i.e., ≥ 1 month; 13). Some alcohol-preferring strains of rodents, such as Indiana P rats [14] and C57BL/6J mice [15, 16, 17], display resistance to quinine adulteration even after a brief alcohol history (i.e., as little as 24 hours). While increased resistance to foot shock-punished self-administration of cocaine has been documented following extended exposure [18, 19, 20, 21], there has been much less work assessing the role of alcohol history in promoting foot shock-resistant alcohol drinking. A study from Radke and colleagues [22] showed that 2 and 4 weeks of exposure to chronic intermittent alcohol produces increased resistance to foot shock punishment during alcohol self-administration compared to mice with self-administration experience only. Together, these findings suggest that alcohol-induced changes may facilitate the development of drinking despite negative outcomes.

While alcohol history may promote drinking despite adverse consequences, the relationship between the two may not be linear (i.e., increased drug exposure does not necessarily mean more likely to be resistant to adverse consequences). For example, in the cocaine literature, a global increase in punishment resistance following extended cocaine access was found to be driven by a subset of animals, while others remained sensitive to punishment [20]. Such individual variation in resistance to punishment has been well-documented, and examination of existing literature on foot shock-resistant drug seeking behavior shows that duration of exposure had no effect on the number of animals within a group classified as shock-resistant [23]. Consistent with this, individual differences in resistance to punishment are evident despite similar alcohol intake at baseline [4, 11, 24, 25, 26]. These studies indicate a role for pre-existing neurobiological differences in driving punishment-resistant alcohol drinking behaviors.

As a whole, these findings raise the possibility that some individuals may demonstrate punishment-resistant drinking regardless of alcohol history, while others may develop resistance to punishment only after a history of alcohol exposure. Thus, the goal of the current study was to expand upon existing literature by examining whether an extensive alcohol history (i.e., alcohol vapor exposure) would alter punished drinking overall, and whether such history would affect individual differences in punished drinking. To do this, male Wistar rats were trained to self-administer alcohol using operant procedures. Rats then underwent foot shock-punished alcohol self-administration sessions, followed by a period of alcohol vapor exposure and subsequent self-administration and punished sessions. We hypothesized that alcohol vapor exposure would promote shock-resistant alcohol self-administration overall, and that this increase would be driven by a subpopulation of rats classified as shock-sensitive prior to alcohol vapor exposure.

## Materials and Methods

### Animals

Male Wistar rats (n=48; Charles River Labs, Wilmington, MA, USA) aged seven weeks upon arrival were used for all experiments. Rats were pair-housed in a temperature- and humidity-controlled vivarium on a 12 hr light/dark cycle (lights on at 7:00 AM). Rats had ad libitum access to food and water for the entirety of experiments, except during the initial self-administration training as noted below. Experiments were conducted during the light cycle. Prior to the start of experiments, rats were handled for at least one min daily for five days. Rats were under continuous care and monitoring by veterinary staff from the Division of Comparative Medicine at UNC Chapel Hill. All procedures were conducted in accordance with the NIH Guide for the Care and Use of Laboratory Animals and institutional guidelines.

### Alcohol self-administration training and baseline

Operant training and testing were conducted in 16 identical operant chambers (31x32x24cm; Med Associates Inc., St. Albans, Vermont, USA) housed inside of sound-attenuating cubicles equipped with exhaust fans. Each chamber contained two levers located on opposite walls. A stimulus light was positioned above each lever and a liquid receptacle was located to the side of each lever. Each completed fixed ratio 2 (FR2) lever response (i.e., 2 lever responses) on the active lever resulted in the delivery of 0.1 mL of solution, as well as the presentation of light and tone cues. Responses on the other lever (i.e., inactive lever) were recorded but produced no programmed consequences. The floor of each chamber was composed of metal grid rods and was connected to an aversive stimulator/scrambler to deliver foot shocks.

Rats were trained to self-administer alcohol using a sucrose fading procedure, similar to [27, 28, 29, 30]. Briefly, rats were water restricted for no more than 24 hr prior to the first sucrose fading session. Rats were then trained to lever press for 10% (w/v) sucrose/2% (v/v) alcohol (10S2A) during a 16 hr overnight session. Subsequent sucrose fading sessions were 30 min and solutions used during each session are as follows: 10S2A, 10S5A, 10S10A, 5S10A, 5S15A, 2S15A, 2S20A, 20A. Rats underwent one session at each concentration with the exception of three sessions at 20A, after which rats were returned to 15A for the remainder of the study. Rats completed 23-28 sessions at 15A before continuing to foot shock-punished alcohol self-administration. The average last 3 sessions of standard (unpunished) self-administration occurring before punished sessions and alcohol vapor exposure are referred to as baseline.

### Foot shock-punished alcohol self-administration

During these sessions, alcohol delivery was paired with foot shock punishment to evaluate drinking despite negative consequences. Punished sessions were identical to standard (unpunished) sessions with the exception that 1 in 4 FR2 responses (i.e., 1 in 8 lever responses) were randomly paired with a foot shock (0.15-0.25 mA, 0.5 sec). Foot shocks occurred at the same time as alcohol delivery and cue (tone and light) presentations. The first FR2 response within a session was never paired with a foot shock. Rats completed 7 total punished sessions: 1 at 0.15 mA and 0.20 mA each and 5 sessions at 0.25 mA. Percent change from baseline was calculated for each rat at each shock session using the following calculation: [(punished alcohol lever responses – average alcohol lever responses of the last three unpunished sessions) / average alcohol lever responses of the last three unpunished sessions] * 100. To categorize rats into phenotypic subgroups, average percent change from baseline for the last 3 sessions at 0.25 mA was determined for all rats. Rats with values in the bottom third (i.e., ≤ 33^rd^ percentile) were classified as shock-sensitive (SS), while rats with values in the top third (i.e., ≥ 67^th^ percentile) were classified as shock-resistant (SR). Rats falling in the middle third were not included in subgroup analyses.

### Alcohol vapor exposure and self-administration

To examine the consequences of extensive alcohol history on SS and SR phenotypes, rats were divided into 2 groups: rats that received chronic intermittent alcohol vapor exposure (n=24) and air-exposed controls (n=24). The first alcohol vapor exposure occurred 72-96 hr after the last punished self-administration session. Alcohol vapor exposure was performed in 4 identical large rat cages (18"L X 9.5"W X 8"H, housing 2 rats each) with custom-built lids housed in a custom-built chamber (La Jolla Alcohol Research, Inc., La Jolla, CA, USA). 95% ethanol was evaporated into fresh air and delivered to cages at a rate of approx. 15 l/min. During each week of vapor exposure, rats were transferred from home cages into vapor chambers for 16 hr per day (5 PM to 9AM) for 4 consecutive days (Monday PM to Friday AM). The air-exposed controls remained undisturbed in home cages.

At the end of each 16 hr vapor session, tail blood samples were collected from 2 rats at 9AM to determine blood alcohol levels (BALs). Blood (approx. 0.2 ml) was collected from unrestrained rats using the tail snip method and centrifuged. Plasma (approx. 0.5 μl) was extracted and assayed for alcohol levels using an AM 1 Alcohol Analyzer (Analox Instruments Ltd, Stourbridge, UK). Rats exposed to alcohol vapor had mean (±SEM) BALs of 163.8±18.5, 164.6±21.1, and 198.4±32.7 mg/dL during the first, second, and third week of exposure, respectively.

As shown in **Figure 1A**, rats underwent the 4-day cycles of alcohol vapor exposure for 3 total weeks, every other week. After each week of vapor exposure, rats underwent 3 standard (unpunished) alcohol self-administration sessions (M, W, F). Following the final self-administration session of the last week, rats completed 5 additional sessions of punished self-administration at 0.25 mA to analyze vapor-induced changes in resistance to foot shock punishment.

**Figure 1.**
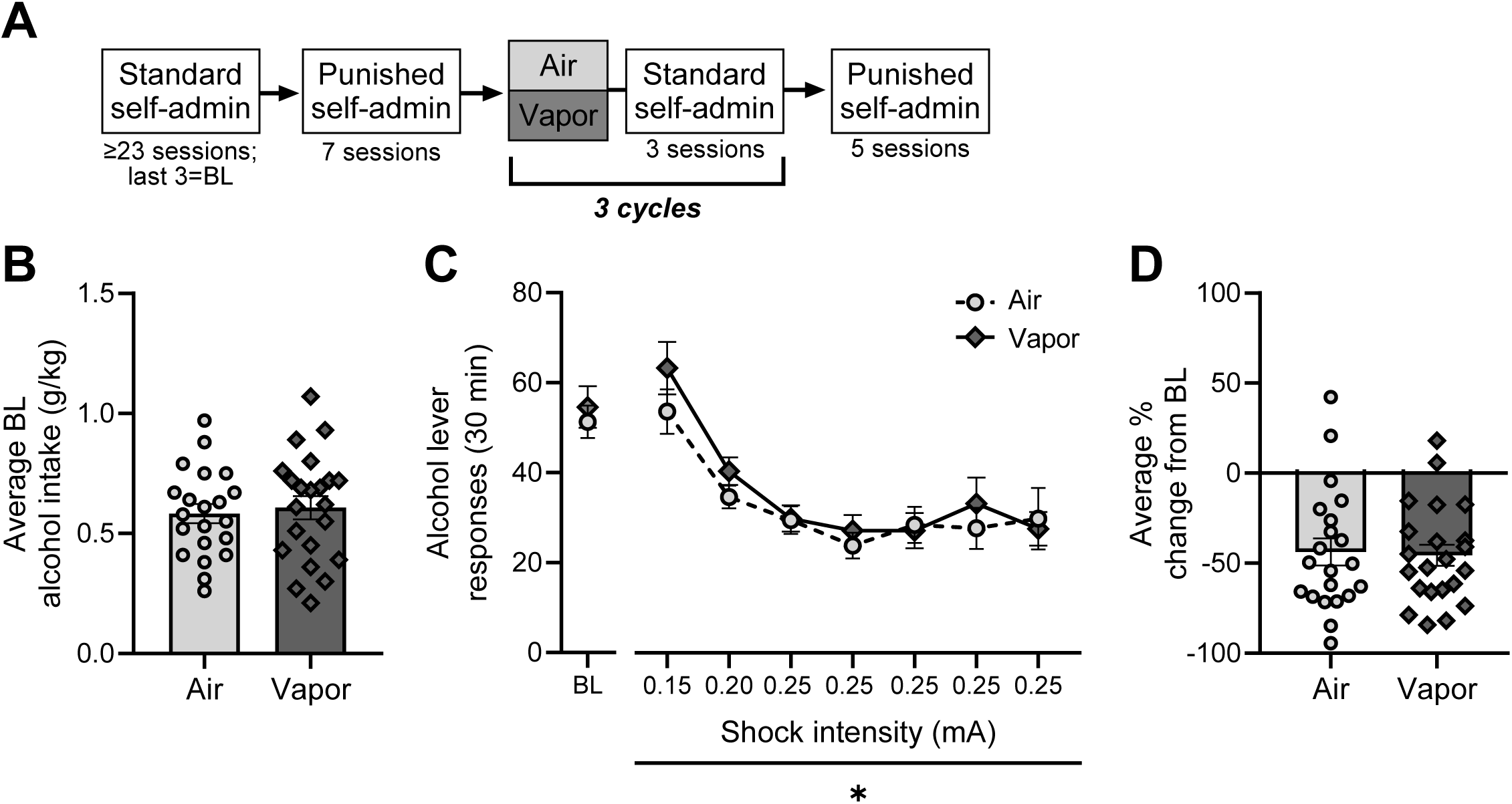
Punished alcohol self-administration prior to alcohol vapor exposure. **(A)** Rats were trained to self-administer 15% alcohol on a FR2 schedule of reinforcement during 30 min sessions, then underwent 7 punished sessions in which 1 in 4 FR2 responses were paired with a 0.5 sec, 0.15-0.25 mA foot shock. Rats then underwent 3 cycles of either air or alcohol vapor exposure followed by 3 sessions of standard (unpunished) self-administration, after which resistance to punishment was re-assessed during 5 punished sessions. **(B)** Average alcohol intake at baseline was similar between air and vapor conditions. **(C)** Air and vapor conditions showed no differences in alcohol lever responses made during punished self-administration sessions before alcohol vapor exposure. Each shock intensity occurs during separate, daily sessions. **(D)** Average percent change from baseline was also similar between air and vapor conditions.

### Statistical analyses

All statistical tests were performed using GraphPad Prism 10.3 (GraphPad Software, San Diego, CA, USA). Average alcohol intake (g/kg; estimated from the number of alcohol reinforcers delivered) at baseline and average percent change from baseline for the pre-vapor punished sessions were compared between air and vapor conditions using a two-tailed unpaired t-test. An ordinary two-way ANOVA was used to examine average alcohol intake (g/kg) at baseline between SS vs. SR subgroups (between-subjects factor) in air vs. vapor conditions (between-subjects factor). Two-way repeated measures (RM) ANOVAs were used to analyze alcohol lever responses and average percent change from baseline between air and alcohol vapor conditions (between-subjects factor) across sessions or weeks (within-subjects factor). Comparisons accounting for both condition (air vs. vapor; between-subjects factor) and subgroup (SS vs. SR; between-subjects factor) were analyzed using three-way RM ANOVAs with session or week as within-subjects factors. Where significant subgroup x session/week interactions were found (and no main effects of condition), data were collapsed across air vs. vapor conditions and analyzed using a two-way RM ANOVA. Post hoc comparisons were performed where appropriate using Tukey’s multiple comparisons test or Fisher’s LSD test. One sample t-tests comparing each group to 0 were used in addition to RM ANOVAs to analyze average percent change from baseline during standard (unpunished) self-administration sessions between subsequent vapor cycles. Three animals from the air condition and two animals from the vapor condition did not meet baseline self-administration criteria of average alcohol intake ≥ 0.20 g/kg and were not included in the study or analyses. Data are presented as mean ± SEM. The threshold for statistical significance was p ≤ 0.05.

## Results

Prior to alcohol vapor exposure, there were no differences between the air and vapor conditions in average alcohol intake (**Figure 1B**) or average alcohol lever responses made during the last 3 standard (unpunished) self-administration sessions (see average baseline alcohol lever responses on left of x-axis break in **Figure 1C**). Examination of alcohol lever responses during pre-vapor punished sessions found a main effect of session [F(3.70, 151.62)=24.40, p<0.0001], indicating decreased responding as shock intensity increased (**Figure 1C**). Importantly, there was no main effect of air vs. vapor condition and no interaction effect, indicating similar punished self-administration prior to vapor exposure. To determine the magnitude of suppression of responding during punished sessions, the average of the alcohol lever responses made during the final three 0.25 mA sessions were normalized to percent change from baseline. Average percent change from baseline was similar between the air and vapor conditions (**Figure 1D**).

### Alcohol vapor exposure induces escalations in alcohol self-administration

We next examined alcohol self-administration following each of the three alcohol vapor exposure cycles. During these sessions, alcohol lever responses were significantly increased in vapor-exposed rats compared to air-exposed controls as indicated by a main effect of vapor condition [F(1,41)=5.62, p=0.023] (**Figure 2A**). There was also a main effect of session [F(5.08, 208.37)=2.87, p=0.015]. There was no significant interaction. To compare the percent change in alcohol lever responses relative to the pre-vapor baseline, we examined the average of the three self-administration sessions after each vapor cycle (**Figure 2B**). Average percent change from baseline was also significantly increased in vapor-exposed rats compared to controls [F(1,41)=6.00, p=0.019]. There was no main effect of post-vapor week and no interaction.

**Figure 2.**
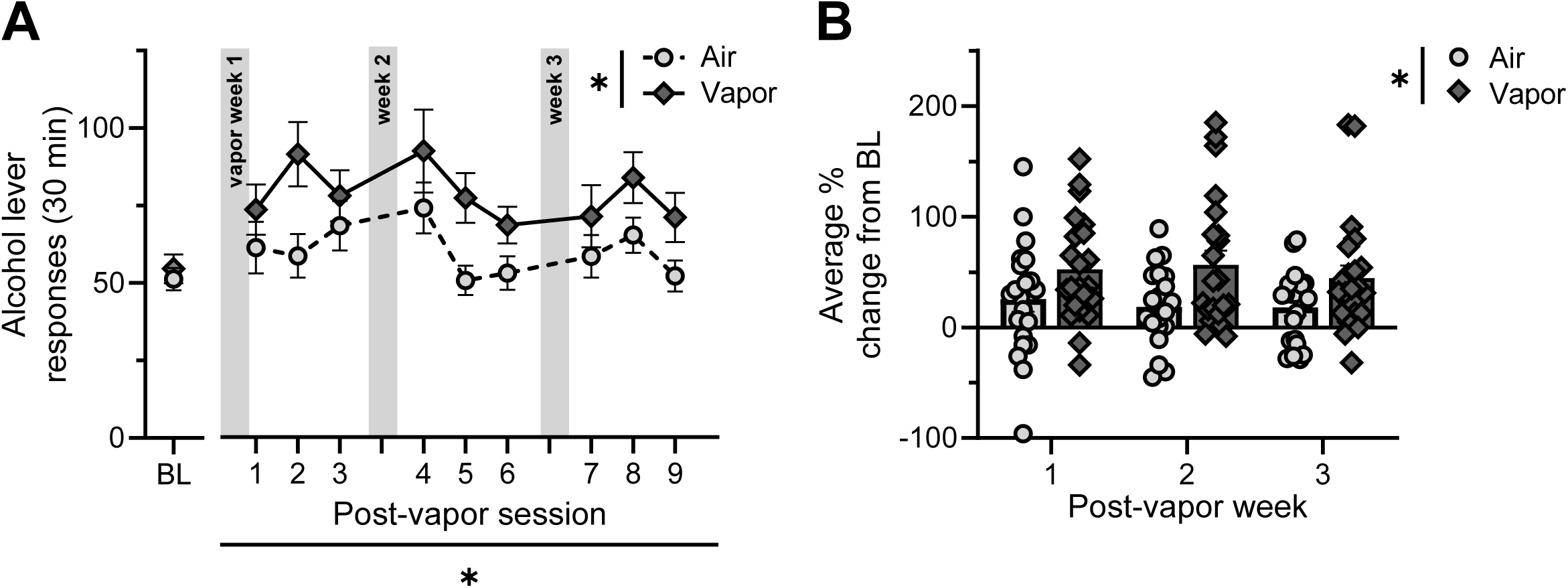
Unpunished self-administration between successive cycles of alcohol vapor exposure. **(A)** Rats in the vapor condition showed a significant escalation in self-administration during unpunished sessions occuring between subsequent cycles of vapor exposure compared to air-exposed controls. **(B)** Average percent change from baseline was significantly increased in vapor-exposed rats compared to rats in the air condition. *p<0.05.

### Alcohol vapor exposure does not affect resistance to foot shock punishment overall

After the three cycles of alcohol vapor exposure, rats underwent 5 punished self-administration sessions to determine whether vapor exposure would affect this behavior, especially given the increased self-administration following vapor exposure observed in Figure 2. **Figure 3A** shows punished self-administration post-vapor (pre-vapor baseline self-administration is shown to the left of the x-axis break). Surprisingly, there was no main effect of vapor condition for alcohol lever responses. There was a main effect of session [F(4, 164)=3.11, p=0.017], indicating decreased responding across the sessions.

**Figure 3.**
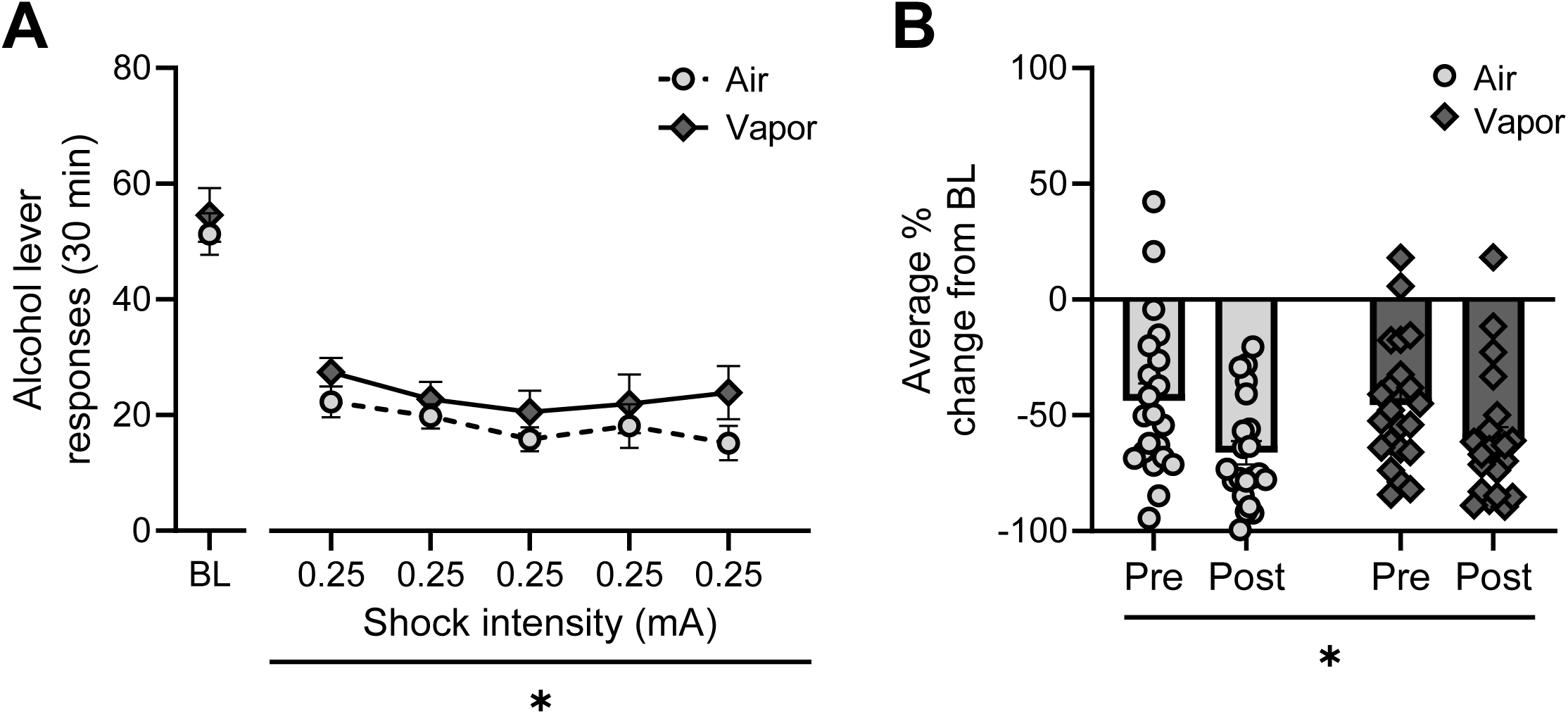
Punished self-administration following alcohol vapor exposure. **(A)** Following alcohol vapor exposure, there were no significant differences in alcohol lever responses made during punished sessions between air and vapor conditions. **(B)** Average percent change from baseline was significantly decreased during post-vapor punished sessions compared to pre- vapor punished sessions regardless of condition. *p<0.05.

We noted that punished self-administration levels were lower post-vapor relative to the initial pre-vapor levels. Therefore, to directly assess suppression of responding, we compared the average percent change from baseline pre- vs. post-vapor via a RM two-way ANOVA (**Figure 3B**; i.e., the average of the final three sessions at 0.25 mA). There was a main effect of time (i.e., pre- vs. post-vapor [F(1,41)=16.93, p=0.0002]), indicating that animals were more sensitive to foot shock punishment post-vapor regardless of air vs. vapor condition.

### Individual differences in resistance to foot shock predict responding in unpunished conditions

We next evaluated whether individual differences in suppression of self-administration during punished sessions before alcohol vapor exposure could predict unpunished self-administration following successive cycles of alcohol vapor exposure. Rats were classified into shock-resistant (SR) and shock-sensitive (SS) subgroups based on average percent change from baseline during the initial (pre-vapor) punished sessions (**Figure 4A**). Rats in the SR and SS subgroups represent the top and bottom thirds of average percent change from baseline respectively; rats falling in the middle third were not included in subgroup analyses.

**Figure 4.**
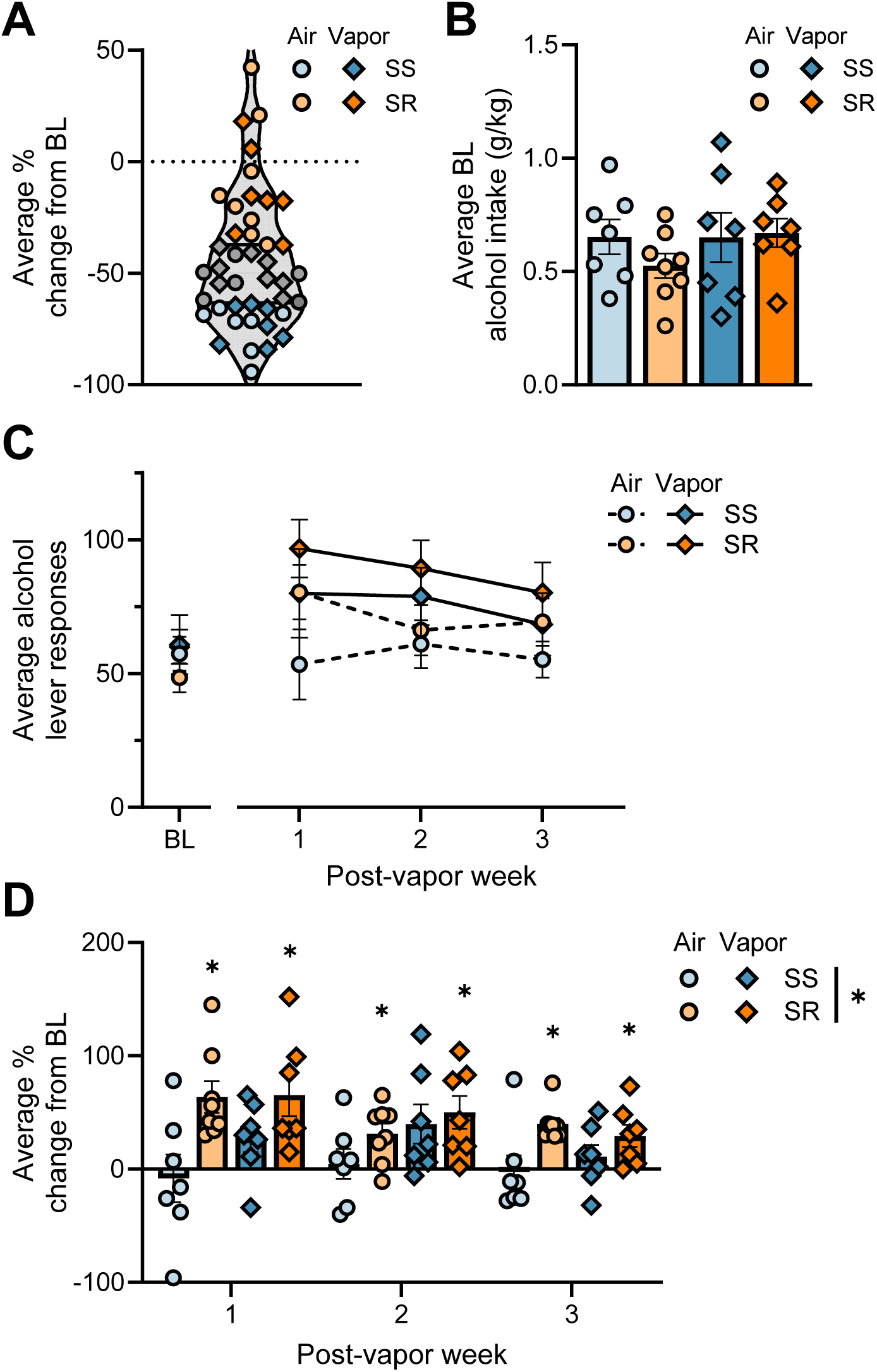
Individual differences in unpunished self-administration between successive cycles of alcohol vapor exposure. **(A)** Individual variability in average percent chance from baseline during pre-vapor punished sessions is apparent when individual data points are plotted. Rats in the top third were classified as shock-resistant (**SR**, in orange) and rats in the bottom third were classified as shock-sensitive (**SS**, in blue). Rats in the middle third (in gray) were not included in subgroup analyses. **(B)** Baseline alcohol intake was not significantly different between subgroups in either air vs. vapor conditions. **(C)** Following each cycle of vapor exposure, rats in the vapor condition showed a trend for increased alcohol lever responses compared to air-exposed controls regardless of subgroup. **(D)** Average percent change from baseline was significantly increased in rats in the SR subgroup. One sample t-tests showed that rats in the SR subgroup significantly increased responding compared to baseline regardless of air vs. vapor condition. SS rats in the vapor condition showed trends for increased percent change from baseline, while no significant change from baseline was observed for SS rats in the air condition. *p<0.05.

Alcohol intake did not significantly differ between subgroups at baseline (**Figure 4B**; see average baseline alcohol lever responses on left of x-axis break in **Figure 4C**). A three-way ANOVA comparing average alcohol lever responses of the three self-administration sessions after each vapor cycle showed trends for a main effect of week [F(1.58, 39.40)=2.87, p=0.080] and vapor condition [F(1, 25)=3.29, p=0.082] (**Figure 4C**). Interestingly, the lack of a vapor effect in this subgroup analysis differs from the escalation in self-administration observed when the entire cohort was analyzed (**Figure 2A**). There was no main effect of subgroup or interaction effects.

Next, we examined average percent change from pre-vapor baseline (**Figure 4D**). A three-way ANOVA showed a main effect of subgroup [F(1, 25)=10.84, p=0.003], with greater increases in responding in the SR subgroup relative to the SS subgroup. There was a trend for a main effect of week [F(1.88, 46.89)=2.71, p=0.080] and no main effect of air vs. vapor condition. There was a trend for a subgroup x week interaction [F(2, 50)=2.91, p=0.064]. To further explore the main effect of subgroup, one sample t-tests were used to compare each group to 0 (i.e., no change from baseline). SR rats in both the air and vapor conditions showed a significant increase in responding relative to baseline following week 1 [air SR: t(7)=4.53, p=0.003; vapor SR: t(6)=3.54, p=0.012] and week 2 [air SR: t(7)=3.49, p=0.010; vapor SR: t(6)=3.45, p=0.014] of vapor. SS rats exposed to vapor showed a trend toward an increase from baseline after week 1 [t(6)=2.22, p=0.069] and week 2 [t(6)=2.31, p=0.061], while self-administration for SS rats in the air condition was unchanged. The significant increase from baseline self-administration rates was maintained for rats in the SR subgroup regardless of condition after week 3 [air SR: t(7)=7.42, p=0.0001; vapor SR: t(6)=3.00, p=0.024]. SS rats in both the air and vapor conditions did not show changes in self-administration following week 3. These findings suggest that SR rats are prone to escalations in alcohol self-administration regardless of alcohol history.

### Individual differences in resistance to foot shock punishment are not altered by alcohol vapor exposure

We next investigated the relationship between punishment resistance pre- vs. post-alcohol vapor exposure. For punished sessions occurring before vapor exposure, a three-way ANOVA comparing alcohol lever responses showed a significant main effect of subgroup [F(1,25)=28.12, p<0.0001] and a significant interaction of subgroup x session [F(4,100)=3.60, p=0.009] (**Figure 5A**). There was no main effect of vapor condition, confirming that subgroups responded similarly prior to undergoing vapor exposure. Post-hoc comparisons demonstrated that SR rats made significantly more alcohol lever responses compared to SS rats for each session (ps≤0.05).

**Figure 5.**
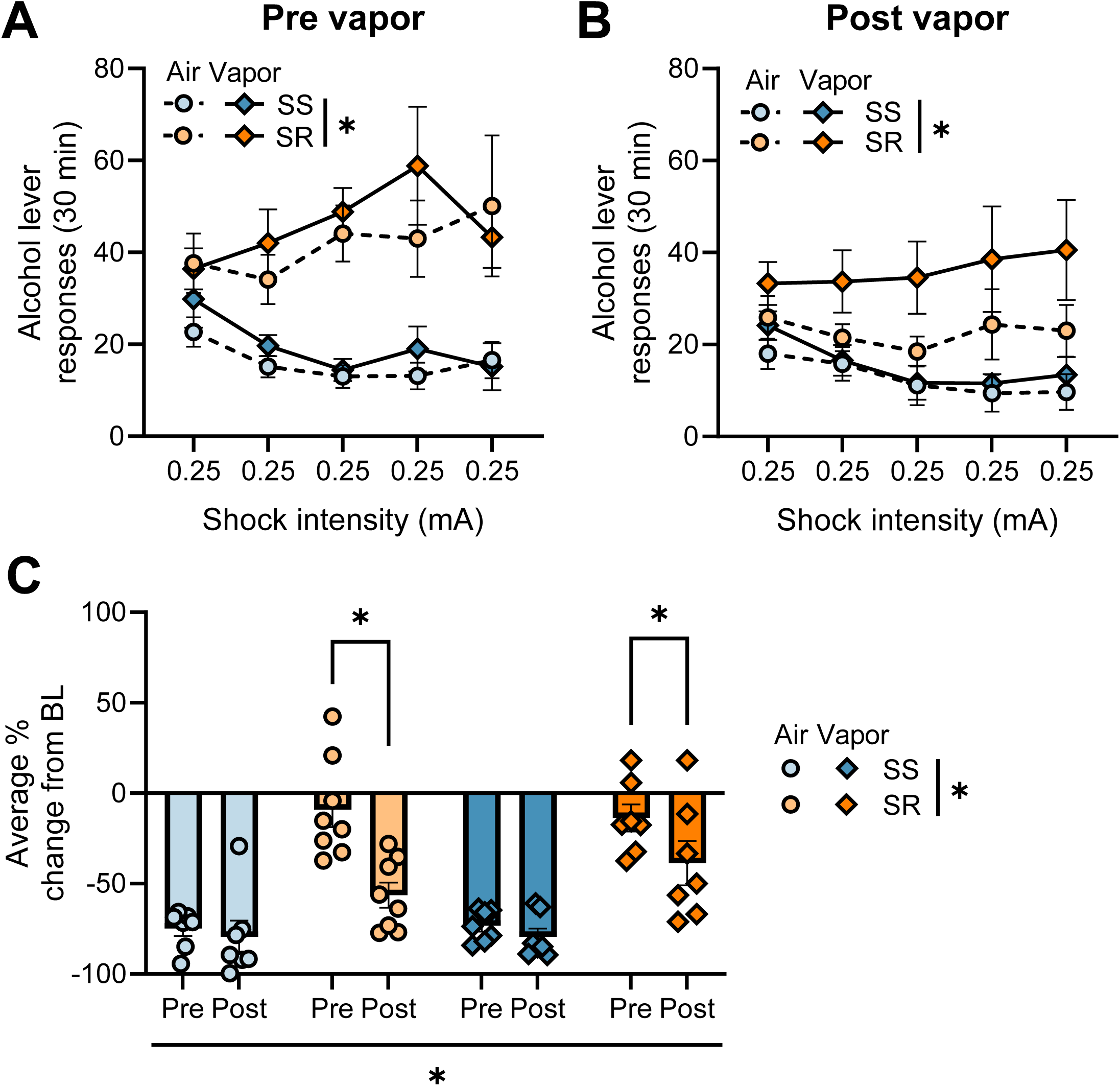
Individual differences in punished self-administration before and after alcohol vapor exposure. **(A)** Rats in the SR subgroup made significantly greater alcohol lever responses during pre-vapor punished sessions compared to the SS subgroup. This significant difference was maintained during post-vapor punished sessions **(B)**. **(C)** Average percent change from baseline was unchanged in the SS subgroup and significantly decreased during post-vapor punished sessions in the SR subgroup. *p<0.05.

After 3 cycles of alcohol vapor exposure, SR rats still showed significantly increased alcohol lever responses compared to SS rats as evidenced by a main effect of subgroup [F(1,25)=11.45, p=0.002] (**Figure 5B**). There were trends for a main effect of vapor exposure [F(1,25)=3.21, p=0.085] and a subgroup x session interaction [F(4,100)=2.07, p=0.090].

Next, to compare the average percent change from baseline for the last 3 punished sessions before and after vapor exposure (**Figure 5C**), we used a three-way ANOVA and found significant main effects of pre- vs. post- vapor [F(1,25)=15.10, p=0.001] and subgroup [F(1,25)=68.59, p<0.0001]. There was no main effect of vapor condition. There was a significant pre- vs. post- vapor x subgroup interaction [F(1,25)=8.41, p=0.008]. Post-hoc comparisons indicated that SR rats showed significantly exacerbated suppression of responding (i.e., decreased average percent change) post-vapor relative to pre-vapor (p<0.0001), while average percent change from baseline was unchanged in SS rats. Together, these data indicate that alcohol vapor exposure has a minimal effect on individual differences in resistance to foot shock punishment.

## Discussion

The present study examined the relationship between alcohol history and drinking despite negative consequences by assessing punished alcohol self-administration before and after alcohol vapor exposure. Two different approaches were used to address this question: analyses using the entire cohort and analyses using the shock-resistant and -sensitive classifications (i.e., top and bottom thirds of the cohort). We hypothesized that vapor exposure would promote increased punished self-administration overall, and specifically in a subset of rats initially classified as shock-sensitive (SS). Unexpectedly, we found that self-administration during punished sessions was significantly lower post-vapor regardless of whether rats had undergone air or vapor exposure. Examination of individual differences in punished self-administration showed that rats initially classified as shock-resistant (SR) significantly decreased responding during post-vapor sessions relative to pre-vapor sessions, while behavior of rats in the SS subgroup was unchanged. We also evaluated unpunished self-administration between successive cycles of vapor exposure. As expected, three cycles of vapor exposure induced increases in self-administration in vapor-exposed rats overall. Interestingly, investigation of individual differences showed that rats in the SR subgroup significantly escalated self-administration regardless of air vs. vapor condition. Together, these results suggest that alcohol history has a minimal impact on punished alcohol self-administration.

Our data show increases in alcohol self-administration following alcohol vapor exposure, consistent with previous findings [13, 31, 32, 33]. However, we found no effect of alcohol vapor exposure on foot shock punished self-administration. This is noteworthy as increased resistance to adulteration of alcohol with quinine following extended alcohol history has been reported [7, 12, 13], although some studies observed no change in resistance [34]. While previous work has suggested that resistance to quinine adulteration of alcohol and foot shock punishment may develop in parallel [25, 35], there are several differences between these paradigms, such as modality (gustatory vs. physical/psychological) and timing (paired with intake vs. paired with seeking) of the negative consequence, which may render them differentially sensitive to prolonged alcohol exposure.

Studies using other paradigms to assess alcohol preference and drinking under conflict after extensive alcohol exposure have yielded mixed results. For example, self-administration of alcohol is resistant to lithium chloride devaluation following chronic intermittent alcohol vapor in mice [36]. In another study employing a modified conditioned place preference task in which mice received foot shock in the alcohol-paired side, chronic intermittent alcohol exposure had no effect on preference for the alcohol-paired chamber [37]. While alcohol vapor exposure has been reported to have no effect on foot shock-punished operant responding for a food reward, it has been demonstrated to increase punishment resistance during alcohol self-administration [22, 38]. Species differences may reconcile these results with those of the present study, as C57BL/6J mice have been documented to require little alcohol exposure before demonstrating drinking despite negative consequences [15, 16, 17]. Together, results from the literature and the present study suggest that resistance to punishment following extensive alcohol exposure may not be a general phenomenon but instead depend on a number of variables, including species, strain, and type and schedule of both rewards and punishment. While a limitation of the present work is that we only used males, sex is another variable which may influence the impact of alcohol history on punished drinking, as female rodents show increased resistance to foot shock punishment and quinine adulteration during drinking tasks [39, 40, 41, 42]. Therefore, it will be important to examine the same behavioral paradigm in female rats.

We observed robust individual variability in punished self-administration behavior. These individual differences were not predicted by alcohol intake at baseline, which is in agreement with previous work [4, 11, 24, 25, 26, 43, 44], although some work suggests that only high-drinking animals develop resistance to quinine [45]. In the present study, sensitivity to punishment was virtually unchanged in the SS subgroup regardless of air vs. vapor condition. This was surprising as we expected a vulnerable minority of rats in the SS subgroup to show enhanced resistance to punishment following alcohol vapor exposure. Indeed, Giuliano et al. [4] reported a subset of rats to develop increased resistance to foot shock punishment during alcohol self-administration, but this was observed after 10 months of alcohol exposure.

In contrast to the unchanged punished self-administration behavior in the SS subgroup, rats in the SR subgroup under both the air and vapor conditions significantly decreased self-administration during post-vapor punished sessions (i.e., displayed decreased resistance to punishment). In addition to the potential procedural-related explanations described above, learning and memory processes may play a role in driving this effect. For example, recent work in rodents and humans suggests that insensitivity to punishment may result from deficits in operant response-outcome learning [44, 46, 47]. While speculative, it is possible that memory consolidation of the punishment contingencies occurred in the time between pre- and post-vapor punished sessions. A related possibility is that foot shock-related memories themselves undergo consolidation to elicit greater aversion [23]. At first glance, this explanation seems unlikely given that the SS subgroup show unchanged resistance to punishment. However, this may be due to a floor effect, as by definition rats in the SS subgroup received much fewer foot shocks than the those in the SR subgroup overall. Regardless, these explanations do not reconcile these results with other work showing that foot shock-resistant alcohol self-administration is a stable trait [4, 25]. More work is needed to determine the stability of punishment-sensitive and -resistant phenotypes.

Given alcohol vapor exposure did not promote increased resistance to punishment in the SR or SS subgroups, the present study indicates that drinking despite negative consequences relies more heavily on intrinsic differences than alcohol history. A caveat of this interpretation is that rats had at least 23 unpunished self-administration sessions before they were phenotyped in the pre-vapor punished sessions, raising the possibility that alcohol-induced changes occur early in exposure history to promote resistance to punishment. However, previous work from Siciliano and colleagues [35] reported that animals who went on to demonstrate resistance to quinine adulteration of alcohol showed unique neuronal activity in medial prefrontal cortex (mPFC) neurons projecting to the periaqueductal gray during their first experience drinking alcohol compared to sensitive animals. Moreover, this differential neuronal response occurred despite having similar alcohol intake at baseline. This suggests that pre-existing neurobiological differences in brain regions such as the mPFC may drive punishment-resistant phenotypes.

Another major finding of this study is that rats in the SR subgroup are prone to escalations in unpunished alcohol self-administration, and this occurs regardless of air vs. vapor condition. This finding is in agreement with previous work showing increased intake of free-access alcohol in rats classified as punishment-resistant [44]. A remaining question is why rats classified as punishment-resistant escalate their drinking following punished sessions despite having comparable intake to their punishment-sensitive counterparts at baseline. It is possible that removal of contingent punishment is sufficient to induce such increases specifically in punishment-resistant rats. A more compelling explanation could be related to differences in motivation. Rats that are resistant to foot shock punishment display higher breakpoints in progressive ratio tasks for alcohol, indicating increased motivation, compared to foot shock-sensitive animals [4, 25]. Moreover, restriction of alcohol access has been demonstrated to lead to escalations in self-administration [48], and the change to every other day self-administration following daily punished sessions in the present study could further promote enhanced motivation for alcohol. Interestingly, we did not observe additive effects of alcohol vapor exposure on self-administration in SR rats, although other studies have shown that increased alcohol exposure also augments motivation for alcohol [49].

In conclusion, the present study demonstrated that alcohol vapor exposure did not facilitate resistance to foot shock punishment during subsequent alcohol self-administration sessions. Moreover, our results suggest that while punishment-resistant animals may be susceptible to increased alcohol intake following punishment, the stability of this behavioral phenotype may depend on a number of conditions. Previous work has begun to uncover the neurobiological mechanisms underlying punishment-resistant and -sensitive phenotypes, finding roles for the medial prefrontal cortex [35], striatum [43, 50], and central amygdala [51], among others. As our findings support a view of punishment resistance as relying on innate differences rather than alcohol-induced underpinnings, future studies may benefit from investigating neurobiological mechanisms underlying vulnerability to develop drinking despite negative consequences.

## Acknowledgements

The authors would like to thank Joseph M. Carew for his contribution to behavioral data collection.

## Author Contributions

**Maya N Bluitt:** Conceptualization, Methodology, Investigation, Data Curation, Formal Analysis, Writing – Original Draft, Writing – Review and Editing, Visualization, Supervision. **Ana Munoz:** Investigation, Data Curation. **Joyce Besheer:** Conceptualization, Resources, Writing – Review and Editing, Supervision, Project Administration, Funding Acquisition.

